# The role of capacity constraints in Convolutional Neural Networks for learning random versus natural data

**DOI:** 10.1101/2022.03.31.486580

**Authors:** Christian Tsvetkov, Gaurav Malhotra, Benjamin D. Evans, Jeffrey S. Bowers

## Abstract

Convolutional neural networks (CNNs) are often described as promising models of human vision, yet they show many differences from human abilities. We focus on a superhuman capacity of top-performing CNNs, namely, their ability to learn very large datasets of random patterns. We verify that human learning on such tasks is extremely limited, even with few stimuli. We argue that the performance difference is due to CNNs’ overcapacity and introduce biologically inspired mechanisms to constrain it, while retaining the good test set generalisation to structured images as characteristic of CNNs. We investigate the efficacy of adding noise to hidden units’ activations, restricting early convolutional layers with a bottleneck, and using a bounded activation function. Internal noise was the most potent intervention and the only one which, by itself, could reduce random data performance in the tested models to chance levels. We also investigated whether networks with biologically inspired capacity constraints show improved generalisation to *out-of-distribution* stimuli, however little benefit was observed. Our results suggest that constraining networks with biologically motivated mechanisms paves the way for closer correspondence between network and human performance, but the few manipulations we have tested are only a small step towards that goal.

## 1. Introduction

Convolutional Neural Networks (CNNs) are extraordinarily successful in a diverse range of complex visual tasks such as object recognition, localisation, image segmentation and captioning (Ren et al., 2017; Chen et al., 2018; Tan & Le, 2021). These successes have inspired research exploring the extent to which CNNs and humans process visual information similarly. For example, previous research has shown that CNNs provide the best account of the representational geometry of object similarity in primate inferotemporal cortex (Khaligh-Razavi & Kriegeskorte, 2014). CNNs also outcompete alternatives in predicting neural firing patterns in various areas of the ventral visual stream by matching hidden layer activity to neural recordings (Yamins et al., 2014; Schrimpf et al., 2018). Others studies have focused on behavioural comparisons, showing some agreement between humans and CNNs in object similarity judgements and shape recognition (Kubilius et al., 2016; Peterson et al., 2017). Such results have generated enthusiasm for the claim that CNNs are the leading model of the visual system (Kubilius et al., 2019; Kriegeskorte, 2015).

However, for all these successes, there remain large gaps between human behaviour and CNN performance. Most often, this disconnect is observed in the form of mistakes of CNNs that are uncharacteristic of human performance, such as differences in how they classify adversarial images (Dujmović et al., 2020) and differences in how humans and CNNs generalise outside their training sets (Recht et al., 2019; Geirhos et al., 2018), with CNNs frequently relying on “shortcuts”, such as texture rather than shape, to classify objects (Geirhos et al., 2019; Malhotra et al., 2020; Geirhos et al., 2020).

There is a second, often overlooked kind of divergence between humans and CNNs — cases in which network performance far surpasses human abilities. Though such high performance is desirable from an engineering perspective, such phenomena pose a challenge for the view that CNNs operate in a similar manner as to humans. In this study, we focus on one such striking example: CNNs that perform well on object classification with datasets of labeled natural images can also learn to classify large datasets of images with random pixel intensities (which to a human observer look like images of TV-static), or natural image datasets with arbitrary mappings between inputs and category labels (Zhang et al., 2017). Indeed, the models even learned to classify ∼ 1 million such random inputs into 1,000 categories, conditions that parallel the `ImageNet` dataset (Deng et al., 2009). These random datasets required only a small multiple of additional presentations (< 5) compared to the unmodified `ImageNet` dataset (Zhang et al., 2017) in order to achieve similar levels of success, with the random pixel images being slightly easier than the shuffled labels. It seems unlikely that humans could do nearly so well under these conditions, especially in the case of the random pixel images.

In this study, we systematically assessed the disconnect between CNNs and human performance on these datasets and explored what conditions can make models perform more like humans. First, we ran a behavioural experiment to confirm that humans are indeed much worse than CNNs in classifying random datasets like those described by Zhang et al. (2017). Next, we investigated why human behaviour diverges so strikingly from CNNs. One possibility is that these differences arise from the vast difference in their visual experiences: While humans have had a lifetime of perceiving structured visual data, CNNs begin training as a ‘blank-slate’. Alternatively, the discrepancy reflects fundamental differences in the processing capacity of CNNs and the human visual system. To explore these ideas we looked at the effect of pre-training CNNs on structured data and compared it to the effect of introducing three biologically relevant capacity constraints – internal noise, bottlenecks and activation functions – to their architectures. We observed that pre-training CNNs on structured data does not reduce their super-human capacity (pre-trained networks in fact became faster at learning random datasets), while adding biologically relevant capacity constraints selectively diminished their ability to learn random data. Finally, we explored whether training models with biologically motivated capacity constraints improved their correspondence with human performance in other respects, specifically, in their ability to generalise to *out-of-distribution* data, such as adding salt-and-pepper or uniform noise to images. However, with minor exceptions, the constrained networks failed to show improvements in generalisation accuracy compared to baseline models. Together, our findings highlight the importance of adding biological constraints to CNNs in order to match human limitations in classifying random data, but also show that additional innovations are required to improve the generalisation capacities of CNNs.

## 2. How well can humans learn random categories?

In previous work, Zhang et al. (2017) trained CNNs on two datasets. One dataset contained images with pixel intensities that were randomly sampled so that each image was akin to a randomly generated vector. These images were then randomly assigned to one of multiple categories. We call this the ‘random pixel condition’. The other dataset consisted of naturalistic images from either the `CIFAR-10` or `ImageNet` dataset, which were randomly assigned to one of the categories at the start of the experiment. Accordingly, images that were members of the same class in the original dataset (e.g., two different images of a plane) could be assigned to different categories, whereas images that belonged to different categories in the original dataset (e.g., an image of a plane and an image of a horse) could be assigned to the same class. We call this condition the ‘shuffled labels’ condition. Zhang et al. (2017) showed that CNNs with different architectures can learn to categorise images from both datasets, even when these datasets contained more than a million images.^1^ The authors also found that CNNs learned to categorise the random pixel dataset more quickly (i.e., fewer training epochs were needed to converge to peak performance) than the shuffled label dataset.

In order to assess how humans learn random data we measured the ability of humans to perform a task similar to that which the networks in Zhang et al. (2017) had been trained on, but limited the training set to far fewer images on the assumption that humans cannot learn to classify over 1 million random images (due to human capacity limitations). This procedure involved categorising either random pixel images or images with shuffled labels. In both conditions the categories lacked any category-defining features.

### 2.1. Methods

We recruited 41 participants using the online platform Prolific (https://www.prolific.co) and randomly assigned each participant to either the random pixel condition or shuffled labels condition. Participants in both conditions completed a categorisation task in which they were asked to learn how to classify 20 images. In each condition, these 20 images were randomly selected from a larger dataset and assigned to one of two categories (10 images per category). Images used in the shuffled labels condition were taken from the Novel Objects and Unusual Names (NOUN) version 2 dataset (Horst & Hout, 2016) and randomly assigned to one of the two categories. This dataset comprises of photographs of inanimate objects which have been judged as being novel using an inter-rater naming consensus (Horst & Hout, 2016) (see Figure 1, dataset II). The novelty of these images ensures that images are unfamiliar to humans. The objects depicted in the images did not share defining features with each other, making this a challenging task to learn (although CNNs can learn an even more challenging task where images with similar features were assigned to different categories).

**Figure 1:**
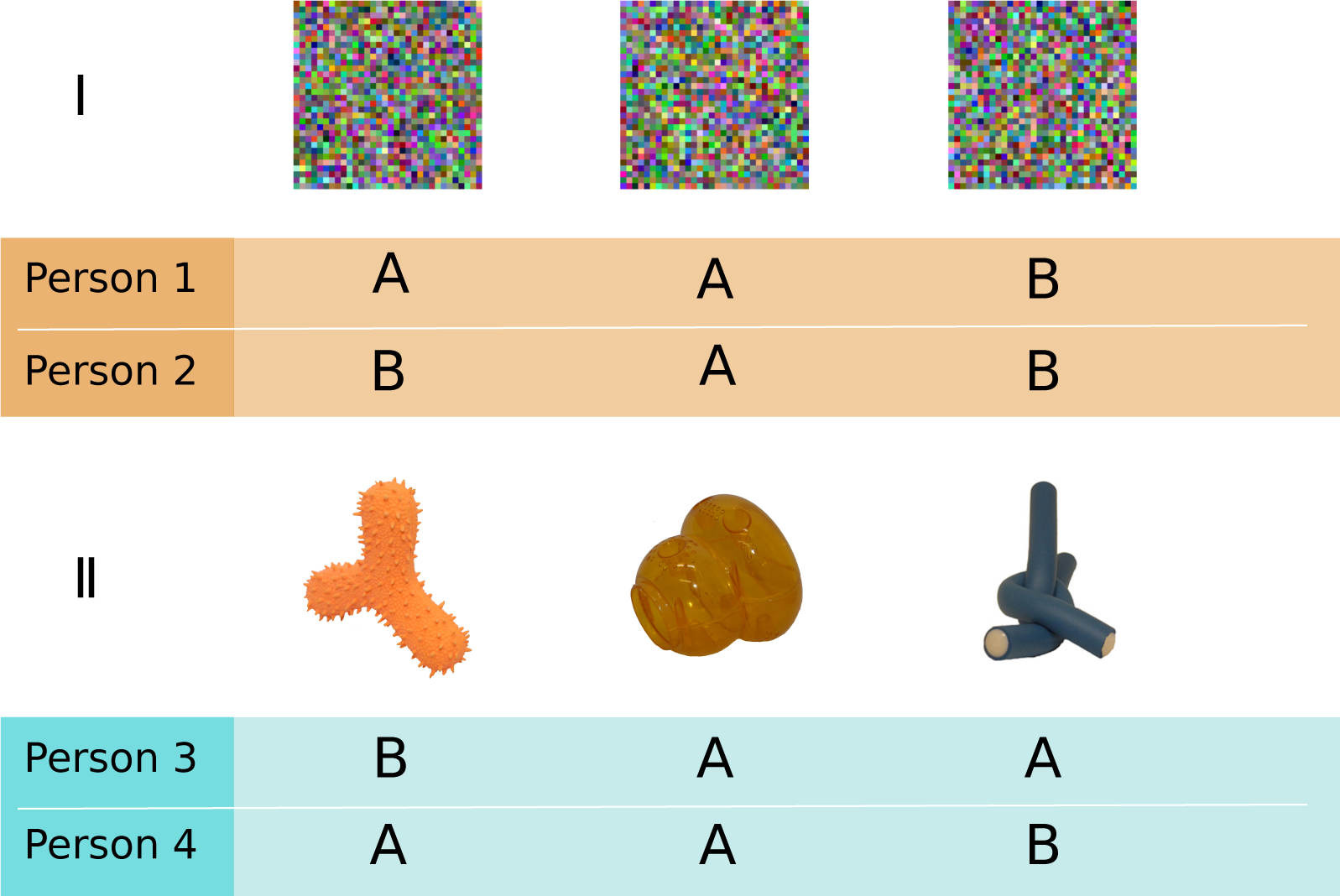
Examples of (I) random pixel images and (II) novel object images shown in the ‘shuffled labels’ condition in the behavioural experiment. Stimuli were randomly assigned to one of two categories, A or B. The stimulus selection and category assignment was different for every participant — e.g., the labels given to Participant 1 were not the same as those given to participant 2.

Images in the random pixel condition were generated by drawing samples from Gaussian distributions for each colour channel, with the mean and standard deviation of each matching the values derived from the `CIFAR-10` dataset (Krizhevsky et al., 2009). The generated images were sized 32 by 32 pixels to match the size of `CIFAR-10` images used in the simulations of Zhang et al. (2017). To overcome any confounding effects of visual acuity of participants, these images were then upscaled (without smoothing) to 224 by 224 pixels. Some examples of these images are shown in Figure 1, dataset I.

Participants were instructed to indicate the categories by pressing either ‘`x`’ or ‘`z`’ on their keyboard. On each trial, a fixation cross was presented on the screen for 250ms. Afterwards, each stimulus was displayed for 300ms. All stimuli were presented on a neutral grey background. Trials were separated by a presentation of an empty neutral grey screen for 500ms. Responses within the first 150ms of presentations were considered too quick and a warning message was displayed to participants to pay more attention to the stimulus before responding. These trials were not considered for analysis. Similarly, we took responses longer than 3000ms as evidence that participants were not attending and these trials were also excluded.

Participants first completed a ‘training’ phase, receiving feedback about their response for each trial. The training phase consisted of 300 trials for the shuffled labels condition and 500 trials for the random pixel images condition. The larger number of trials for the random pixel images condition was motivated by a pilot study that showed participants struggled to learn this task and we wanted to ensure that they received sufficient training. Breaks of up to 5 minutes were included midway through, and upon completion of the training trials. Participants then completed a ‘test’ phase, in which they no longer received any feedback, with the number of trials equal to 20% of training trials (60 and 100 trials, respectively).

### 2.2. Results and Discussion

We computed the average categorisation accuracy of participants for the second half of the training phase and for the test phase (Figure 2). In the test phase of the random pixel condition, participants’ accuracy (M = 0.529, SD = 0.11) was not statistically different from chance (One-sample *t*-test, two-tailed, *t*(20) = 1.229, *p* = 0.233). In contrast, most subjects learned to categorise the images in the shuffled labels condition (M = 0.895, SD = 0.157). The difference in accuracy between the two groups was statistically significant (two-tailed, *t*(40) = 8.639, *p <* 0.001, Cohen’s d = 2.699, 95%CI=[1.825; 3.572]).

**Figure 2:**
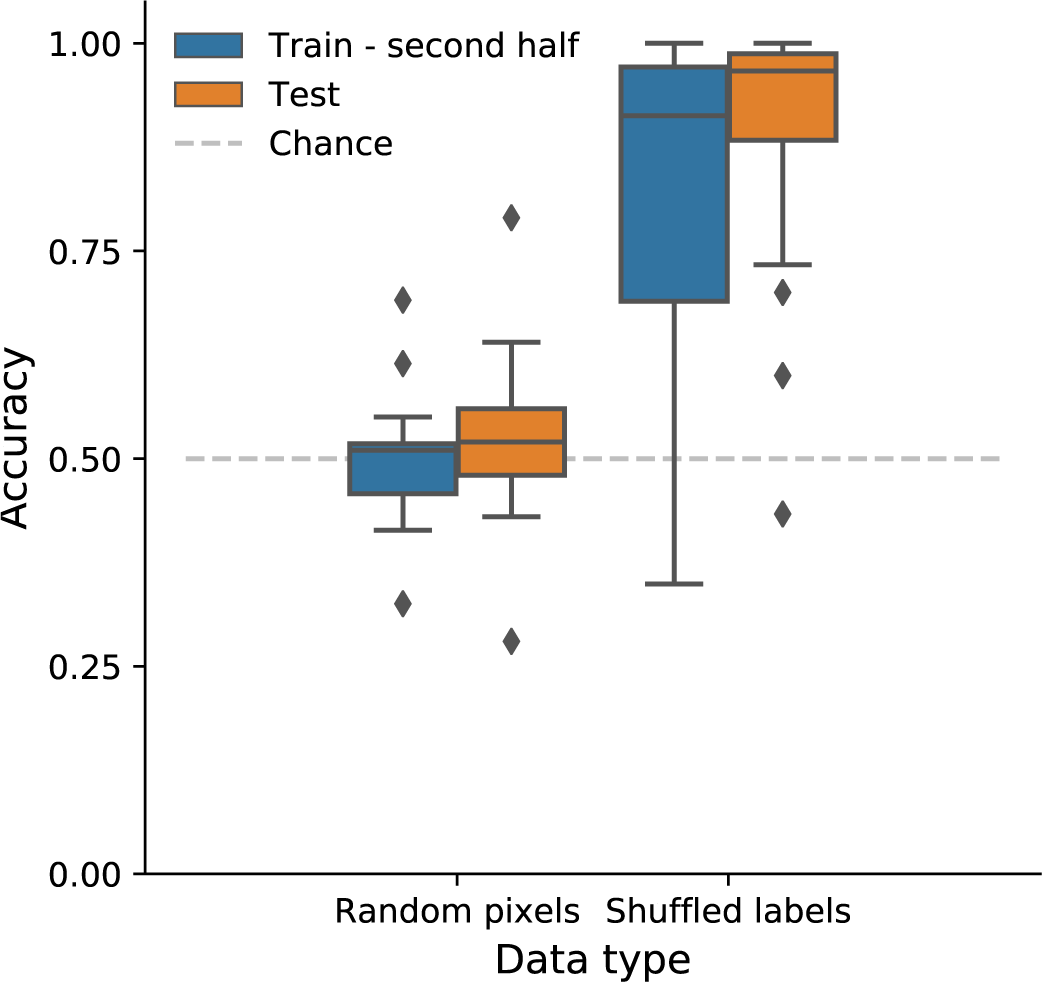
Humans do not improve much over chance levels when classifying random pixel images. Boxes show average accuracy for the second half of the training phase (blue) and the test phase (orange)

The results of the behavioural experiments match our expectations. Participants were near or at chance performance in categorising the 20 random pixel images into two categories, even after 500 learning trials. Three individuals who succeeded in achieving above-chance performance were invited to answer a post-experiment debrief question about what strategy they used to complete the task. Out of three participants, two replied, specifying they tried to distinguish stimuli by focusing only on certain patches within the images (such as the top-left quadrant) and paying attention to the local properties of that region (e.g. “This one had more red overall”), rather than paying attention to individual pixels. Although these heuristics allowed some participants to achieve an above chance performance in a 2-way categorisation task involving a small dataset of 20 images, they would become much more difficult to execute when increasing the number of examples or categories. Scaling the strategy to a 1,000-way classification of ∼ 1 million images is infeasible.

In contrast to the random pixel condition, almost all participants were able to learn to classify the 20 novel objects into two categories in the shuffled labels condition with over 80% accuracy in the test phase. It is unlikely humans can rival the capacity of a network to learn to classify ∼ 1 million images into 1,000 random categories, but the results showed another disconnect between humans and neural networks — humans find it more difficult to learn to classify random pixel images than novel objects with shuffled labels, whereas Zhang et al. (2017) observed that models did not show any preference for learning the structured data (shuffled labels) compared to random pixels. Indeed, CNNs took *longer* to learn to categorise stimuli in the shuffled labels condition compared with the random pixel condition. Together, these results highlight a fundamental misalignment in information processing between the models and the human visual system.

## 3. Improving correspondence between networks and human behaviour

The experiment in Section 2 established two types of differences between humans and CNNs, namely, a) humans struggle to learn even a small set of unstructured data (random pixels condition), while CNNs are much better at learning this type of data, and b) humans are better at learning structured data that is randomly assigned to categories (shuffled labels condition) while CNNs are equally capable of learning both naturalistic images with shuffled labels and unnatural images with random pixels (Zhang et al., 2017).

Here we explore whether different training environments and architectural modifications can make CNN performance more qualitatively similar to the human results. First, in an attempt to model the effect of human experience — participants spend a lifetime classifying naturalistic images prior to taking part in the experiment — we investigated whether pre-training CNNs on datasets of naturalistic images limits the learnability of the unstructured datasets. Second, we introduced various biologically inspired constraints to CNNs designed to reduce network capacity to learn unstructured data, namely, the introduction of processing bottlenecks, internal noise, and bounded activation functions, as detailed below.

### 3.1. The role of experience

The divergence in learning and memorisation abilities between humans and CNNs might be explained by the differences in their visual experience. Living in a world of statistical regularities and predictable structures, human beings learn and adapt to these reliable features. One consequence of this could be becoming less adept at dealing with information that lacks structure. For example, experts in a given domain like chess are remarkably better than novices in remembering and reconstructing chessboard configurations from games, but perform on an equal footing with less experienced players when reconstructing random configurations of pieces (Chase & Simon, 1973). Indeed, the effects of early visual experience on development and behaviour can be profound. For example, limiting the visual experience of kittens exclusively to either horizontal or vertical bars makes the animals insensitive to the oriented stimuli that they were not previously exposed to (Blakemore & Cooper, 1970). We examine whether a comparable argument can be extended to CNNs. Does experience with the natural world, operationalised as pre-training a network on a dataset of natural image, set limits on what kind of data can be further learned by that network? Would this experience cause learning unstructured data, such as random pixel images, to be more difficult?

#### 3.1.1. Methods

We trained `ResNet-50`, a standard CNN architecture (He et al., 2016), to classify either the shuffled label `CIFAR-10` dataset or random pixels images.^2^ In order to assess the impact of past experience with structured data, we compared a baseline condition in which the weights were randomly initialised (using uniform Xavier initialisation (Glorot & Bengio, 2010)) to CNNs with the weights learned by pre-training the network on the `ImageNet` dataset (Deng et al., 2009), a large database of annotated naturalistic images. For the pre-trained networks we included two training conditions. In a transfer learning condition (Yosinski et al., 2014), we ‘froze’ the weights of all convolutional and pooling layers (meaning that these weights were not updated during further training) and re-initialised the weights of the fully connected layers. In a ‘fine-tuning’ condition we allowed all the weights of the network to be modified during training, similar to the baseline condition. We used three different seeds for every condition. We trained the networks until convergence once for every condition. As every network converged in under 70 epochs, we used this as a boundary for the other two instances. Training hyperparameters used in the simulations are listed in Appendix F, Table F.1. The model architecture was modified by inserting one hidden layer with 1024 units

immediately prior to the output layer, adding ∼ 2 million parameters to the network. This modification was necessary because initial tests indicated that without it, when all convolutional layers in the network were frozen, none of the datasets could converge to sufficiently high accuracy.

Images from `CIFAR-10` were upsampled with smoothing (Burt & Adelson, 1983) from 32 × 32 pixels to 224 × 244 pixels to match the input size of the pretrained networks. Random pixel images were created in resolution 32 × 32 pixels, then resized without smoothing to 224 × 224, forming a grid of 32 squares of size 7 × 7. All images were represented in RGB format. The size of the random pixel dataset matched the size of the `CIFAR-10` training data at 50,000 images. The datasets were the same in each iteration for all models.

#### 3.1.2. Results and Discussion

We assessed the accuracy of the networks on the unmodified data, random pixel and shuffled labels datasets in the three training conditions (Figure 3). Strikingly, pre-training the networks on `ImageNet` did not prevent them from learning the random datasets. Both random pixels images and shuffled labels data were eventually learned with perfect accuracy across all conditions.^3^ While pre-training accelerated learning on the structured (`CIFAR-10`) dataset, it also made learning faster on the unstructured datasets, for both the fine-tuning and transfer learning conditions. Crucially, we did not observe an interaction between pre-training and the type of data being learned. That is, we did not find pre-training to be preferentially beneficial to the speed of learning the `CIFAR-10` dataset compared to the random pixels dataset.

**Figure 3:**
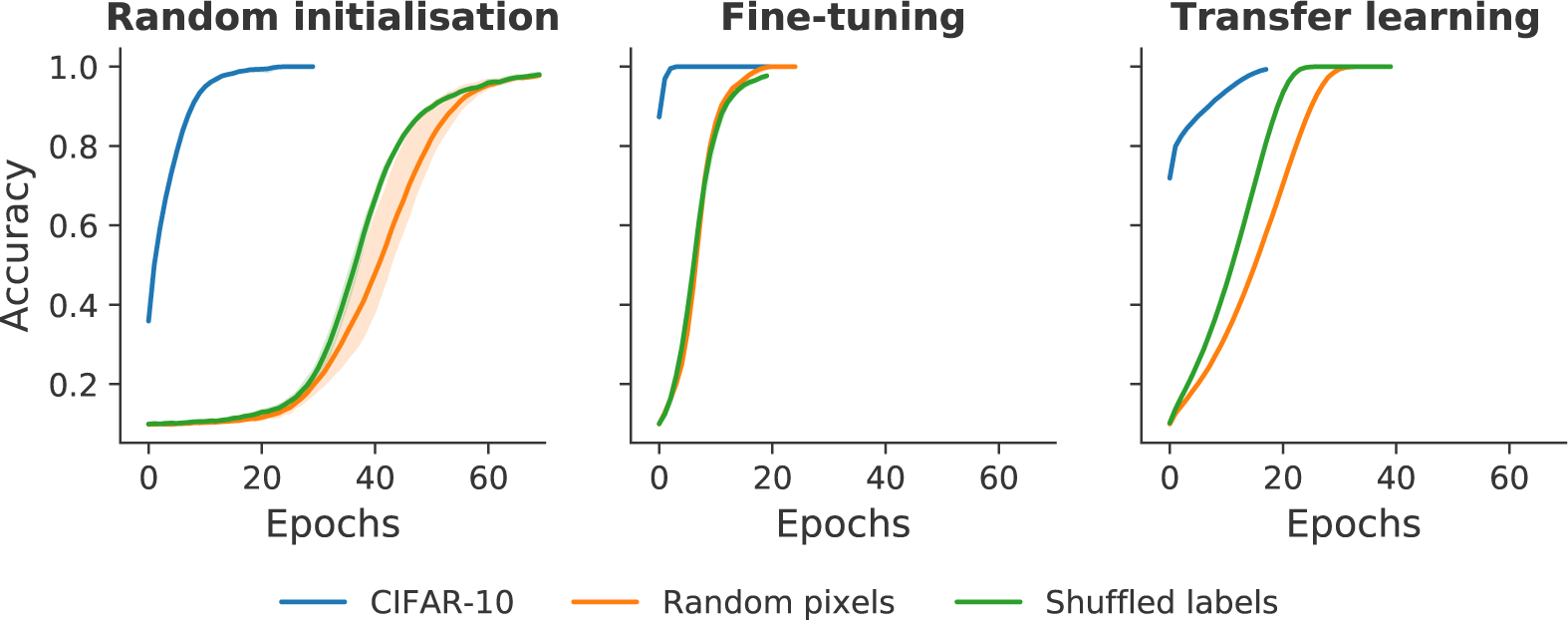
Pre-training on `ImageNet` does not prevent learning random data. Training accuracy is shown for `ResNet-50` networks trained to classify either unmodified `CIFAR-10` (blue), the same dataset with shuffled labels (green), or Random pixel images (orange). The models were either trained from a random initialisation (**Left**), or were pre-trained on `ImageNet`, fine-tuning the entire network (**Right**), or transfer learning: freezing the convolutional weights and training only the classifier (**Center**).

These results do not corroborate the hypothesis that experience with natural images restricts the network’s ability to learn random data. Indeed, the networks pre-trained with `ImageNet` learned the random dataset faster than those with randomly initialised weights.

### 3.2. The role of biological constraints

Next, we investigated whether introducing biologically-inspired constraints would limit the capacity of CNNs to learn random data. We experimented with three such constraints (see Figure 4).

**Figure 4:**
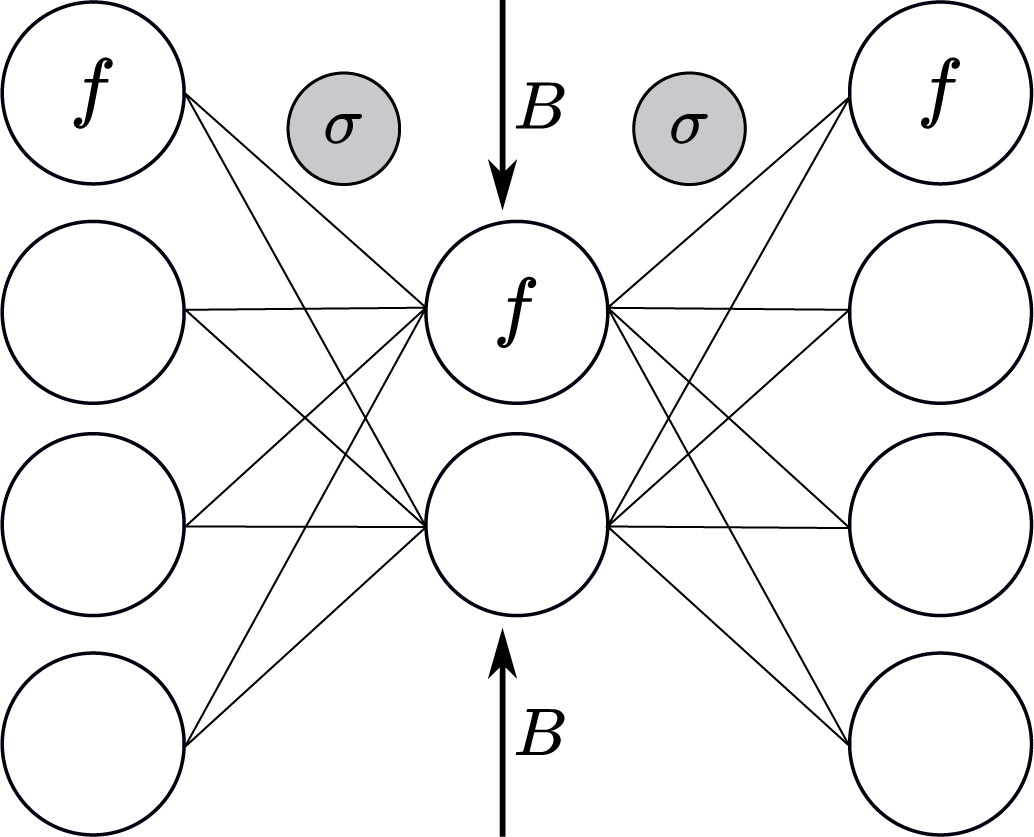
Illustration of biologically-motivated capacity constraints. Gaussian noise (σ) was added to the output of activation functions of hidden units. Hidden units had Sigmoid instead of ReLU activation function (*f*). The bottleneck (*B*) was introduced by reducing the number of convolutional filters in a layer, modifying the architecture of the network.

First, we introduced **internal noise** — or variability — in the activations of units in the CNN. There are various sources of internal noise in the brain. For example, it is a long-standing observation that presentation of the same stimulus can often result in different neural response patterns (Stein et al., 2005). This variability can be observed at multiple spatial and temporal scales, caused by environmental factors or specific cell properties. The largest source of neuronal noise is synaptic, produced by stochasticity in neurotransmitter release, which can have a substantial net effect on the behaviour of post-synaptic cells (Stein et al., 2005). Adding internal noise to the activation values of hidden units in a neural network reduces the precision of the signal, and in turn, may reduce the capacity of the network to encode large numbers of random categories. If the precision of network computations is more critical for the random datasets due to the networks having to rely on fine-grained idiosyncrasies in the data, then internal noise should start to drown this delicate signal and selectively limit CNNs’ ability to classify random data.

Another biologically inspired concept we explored is a **bottleneck** in a hidden layer. Bottlenecks could, in principle, play a pivotal role in reducing representational complexity to allow efficient processing in the perceptual system (Essen et al., 1991) by discarding irrelevant details and passing on the most reliable and robust features of the inputs (Evans et al., 2022). We hypothesised that if category members share some regular features — as is the case with `CIFAR-10` categories — a bottleneck should encourage these features to be discovered, leading to robust representations. On the other hand, if no common features for each category can be detected, then a bottleneck is expected to have a large impact on network performance.

Lastly, we considered the role of the **activation function** on a neural network’s capacity. Typically, modern deep learning models use activation functions which permit a large range of activations. For example, the rectified linear (ReLU) units lead to output activations that are bounded at the lower end but unbounded at the upper end. In contrast, the Sigmoid activation function, which is often encountered in connectionist models of cognitive processes and neurophysiological models of neural population dynamics (Wilson & Cowan, 1972), bounds output activation at both the lower as well as the upper end. Although there is lack of consensus about the biological plausibility of Sigmoid versus ReLu activation functions (see Glorot et al., 2011), we focus on the effect of their range on regulating the capacity for representing information. The activation of ReLU units can grow as large as needed, and their representational capacity is only limited by the degree of precision imposed by the numeric data. Units using the Sigmoid activation function, on the other hand, have their output values limited within the range (0, 1). When paired with the limited precision caused by floating-point operations in computers, they therefore provide a natural constraint on the representational capacity of these units.

#### 3.2.1. Methods

We used two model architectures, `small-inception` and `small-alexnet`, adapted from Zhang et al. (2017, p. 12). These models were not pre-trained as in Section 3.1, but started with randomly initialised weights. While the investigation into the role of experience demanded working with models pre-trained on a large natural image dataset, the architectures selected for this study are scaled down versions of popular models, reducing computational demands. This reduction in trainable parameters permits a wider and more thorough exploration of relevant conditions and hyperparameters.^4^

We implemented internal noise by adding random values drawn from a Gaussian distribution (*µ* = 0, σ = [0 : *x*]) to the output of the activation function of every hidden unit in the neural network.^5^ We varied the standard deviation of internal noise (σ) from 0 − 1.2, with an increment of 0.05 for networks using ReLU activation units, and from 0 − 0.2, in 0.02 steps for Sigmoid networks.

Inspired by Lindsey et al. (2019), we implemented a bottleneck by reducing the number of convolutional filters in the first convolutional layer. This procedure aims to model the reduction from photoreceptors to ganglion cells in the retina. We experimented with several values for the number of filters in the bottleneck and settled on the smallest number possible without drastically impairing performance on the `CIFAR-10` validation data. As a result, we reduced the number of filters in `small-inception` from 96 → 2 and from 200 → 8 for `small-alexnet`.^6^ Although this manipulation only reduces the total number of trainable parameters by a small amount, it has been demonstrated to qualitatively alter the types of filters learned in early convolutional layers, resulting in the development of center-surround receptive fields in the first layer and a prevalence of Gabor-like filters in subsequent layers, similar to what is observed in the human visual system (Lindsey et al., 2019).

We generated model instances for all four combinations of the activation (ReLU or Sigmoid) and bottleneck (yes or no) constraints. Each network instance was trained to classify the three types of datasets described in Section 3.1: (a) the unmodified `CIFAR-10` dataset, (b) a version of the `CIFAR-10` dataset with shuffled labels (c) random pixel images. Like the simulations in Section 3.1, we trained the networks until convergence. Both networks converged under 100 epochs for all data types. Subsequently we used 100 epochs as a boundary in further runs. We used a batch size of 128 (39,000 steps).^7^ We used learning rates which lead to the fastest convergence on all datasets (determined empirically; *lr* = 0.1 for `small-inception` and *lr* = 0.01 for `small-alexnet`). A step decay algorithm was used to decrease learning rate by 5% after each epoch. Further information about the hyperparameters used in these simulations can be obtained in Appendix F, Table F.1. Internal noise was added to the output of the activation function of each hidden unit in every convolutional or fully-connected layer of the CNNs. All results were averaged over four different random seeds for each network instance, with new random pixel images and a shuffled labels permutation for every seed. The same hyperparameters were used for all datasets and seeds.

#### 3.2.2. Results and Discussion

Results from all simulations are summarised in Figure 5 (top row), which shows accuracy on the training set after 100 training epochs for all three datasets for the `small-inception` architecture. Each panel plots the accuracy after 100 epochs as a function of the internal noise. Performance on the `CIFAR-10`, random pixel and shuffled labels datasets are shown using blue, orange and green lines. Bold lines show performance without a bottleneck while dashed lines show performance using a bottleneck.^8^

**Figure 5:**
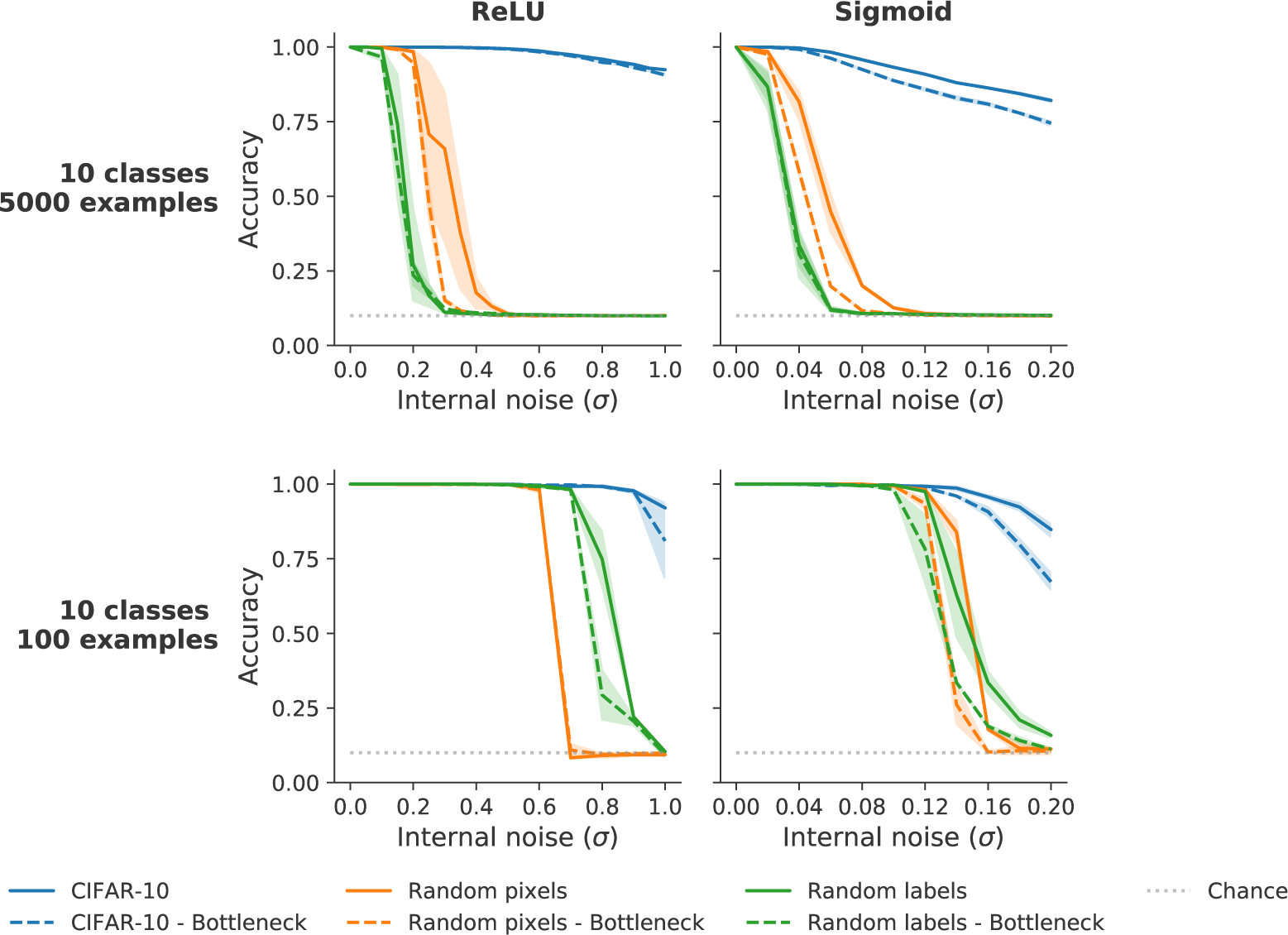
**Top:** Training with biological constraints severely reduces CNNs’ ability to learn random datasets. These figures represent the results described in Section 3.2. **Left**: ReLU activation function. **Right**: Sigmoid activation function. The graphs show the impact of the capacity manipulations on training accuracy (*y*-axis) for the three types of datasets (blue: unmodified `CIFAR-10`; green: modified `CIFAR-10` with shuffled labels; orange: dataset of random pixels images). The internal noise levels used are plotted on the *x*-axis. Solid lines indicate model accuracy for models without a bottleneck; dashed lines indicate models with a bottleneck. **Bottom**: CNNs trained on smaller datasets require greater values of internal noise for random dataset learning to decrease. The figures represent the results described in Section 3.3

We observed that internal noise had a large, selective effect on the ability of the network to learn the two unstructured datasets. That is, within the explored range of values, noise had little impact on the training set classification accuracy for `CIFAR-10` images (although the effect was more noticeable for `small-alexnet` networks). On the other hand, increasing noise values progressively hindered performance on the random pixel and shuffled labels conditions, with training accuracy quickly dropping to chance. Comparing networks using ReLU hidden units with those using Sigmoid units, it appears changing the activation function, by itself, did not impact the ability of the networks to learn to classify the random datasets. However, the activation function interacted with the networks’ tolerance to internal noise, with ReLU unit networks requiring ∼ 5 times more noise compared with Sigmoid units to produce a similar decline in accuracy for the random pixels and shuffled labels conditions (see different scales on *x*-axes). Nevertheless, the same pattern of results was obtained with the two activation functions, with a selective effect of increased noise on the two unstructured datasets but little impact on classifying `CIFAR-10` images.

We also observed that the bottleneck manipulation was not sufficient to prevent learning of unstructured data on it’s own, although its effect interacted with the degree of internal noise and the activation function. That is, for networks with a bottleneck, a slightly lower internal noise value was needed to prevent the network from learning unstructured data.

### 3.3. Does noise affect capacity?

The findings in Section 3.2 suggest that adding internal noise to the activations of hidden units was the most effective method for reducing CNNs’ ability to learn random datasets. The next step was to work out *why* it was effective. Our hypothesis was that this is because noise decreases capacity. If this is the case, then even after inserting noise the network should be able to learn the random pixel and shuffled labels datasets, as long as they do not exceed the now lowered capacity. We manipulated this demand on the network by decreasing the number of exemplars in each category.

We trained the same model architectures used in Section 3.2 on identical tasks — classifying `CIFAR-10` images with either unmodified or shuffled labels, as well as random pixel images. However, we reduced the size of the datasets so that each class contains fewer examples than the full dataset. Lowering the size of the dataset lowers the difficulty of the problem the network has to solve because the model has to fit fewer data points. We hypothesised that if adding internal noise affects capacity, then by reducing the dataset size, and therefore task difficulty, the level of internal noise necessary to suppress random data learning would increase. For example, if CNNs with internal noise σ *_f ull_* were no longer able to learn a dataset of 50,000 random pixel images, they could still learn a smaller dataset of, e.g., 1,000 examples. That is, a larger noise value, σ*_part_* (σ*_part_ >* σ *_f ull_*) would be required to prevent the CNNs from learning the smaller dataset.

#### 3.3.1. Method

We recreated the simulations in Section 3.2, using the same network architectures, but decreased the number of classes and examples per class in each dataset. For the unmodified `CIFAR-10` and the shuffled labels conditions, a stratified sampling procedure was used to ensure an equal number of examples were sampled from each class. For the random pixels condition, an appropriate number of images were generated and randomly assigned to each class. The tested datasets consisted of 10 classes, with 100 examples per class.

Training the neural networks on the smaller dataset with the same batch size and number of training epochs used when training on the full dataset would result in overall fewer weight updates. Therefore, we scaled the number of training epochs and the batch size, so that networks learning the smaller dataset would have the same number of training steps as those trained on the full dataset. See Appendix F for specific values.

#### 3.3.2. Results and Discussion

The pattern of results corroborates our hypothesis, with greater values of internal noise needed to prevent CNNs from learning the unstructured datasets. This is easily seen when compared with results from training on the datasets of 50,000 images used in Simulation 3.2 on the top row of (Figure 5).^9^

There was another interesting, unanticipated effect of training the models with smaller datasets, namely, the relative difficulty of learning random pixels versus shuffled labels swapped when trained on 100 compared to 5,000 images per category in the ReLU condition. However, we do not consider this finding to be robust, as the results were less clear in the Sigmoid condition. We also found that the relative difficulty of the two random conditions varied in the pre-training simulationss in Section 3.1, influenced by various factors such as hyperparameter choice. We do not have a good explanation for this effect, but a successful model of human vision should not only explain the very limited capacity of humans to learn unstructured data, but also the much greater difficulty humans experience in learning the random pixel stimuli. Although our introduction of biological constraints is a step in the right direction, we have not identified a condition that allows to disassociate the ability to learn these two forms of random data within the existing CNN framework.

## 4. Do capacity constraints improve generalisation?

In the simulations in Sections 3.2 and 3.3 we have shown how adding biologically motivated constraints makes CNNs more similar to humans in that the neural networks’ capacity to learn random data is reduced without a notable disruption to their ability to classify natural images. We now consider whether learning with these constraints has consequences for other areas where correspondence between CNN and human performance is lacking, namely, out-of-distribution (*o.o.d.*) generalisation. That is, CNNs struggle to generalise to images that differ in their statistics from images that the network was trained on (Geirhos et al., 2018; Sinz et al., 2019). Here, we look at the performance of CNNs when presented with images that have been manipulated with image distortions not observed during training. This problem is challenging for CNNs, with performance of models degrading rapidly after even small amounts of image distortions or other perturbations (Geirhos et al., 2019). Human performance also deteriorates under these conditions, but more smoothly and to a much lesser extent (Geirhos et al., 2019). We tested whether training CNNs with resource capacities limited by biologically inspired constraints on naturalistic data will force the networks to learn more general representations that support improved *o.o.d.* generalisation.

### 4.1. Methods

To test *o.o.d.* generalisation, we followed the method used by Geirhos et al. (2019) to modify the `CIFAR-10` test set. We created two test sets, each of which converted images from the `CIFAR-10` test set to grayscale and added a type of noise that was not seen during training. In the first set, we added uniform (zero-centred) noise to each image — that is, we modified each pixel by a random value drawn from the distribution [−*a*, +*a*], in which we varied the value *a* in the range [0, 0.5] in increments of 0.05. In the second dataset, we applied salt-and-pepper noise to each image, where each pixel of an image can be switched to black or white with a defined probability, *p*, in the range [0, 0.5], varied in increments of 0.05 (5%). Examples of both types of image degradation are shown in Figure 6.

**Figure 6:**
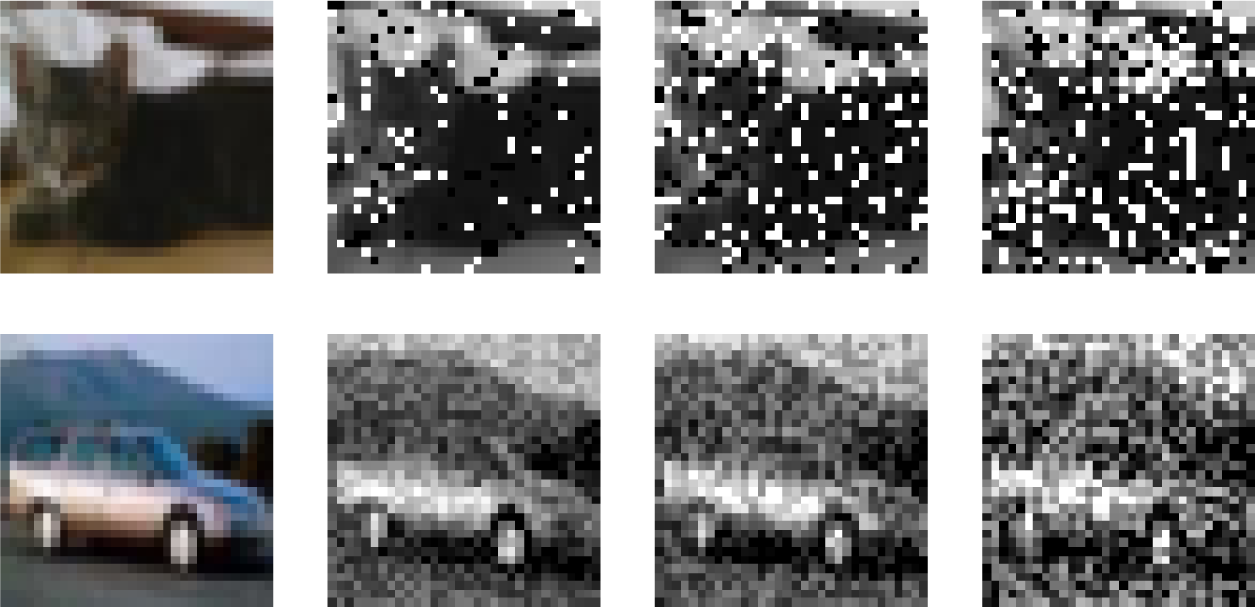
Example CIFAR-10 images with different amounts of salt and pepper noise (top) or uniform noise (bottom).

We trained the same set of CNN architectures as in Section 3.2, using the same hyperparameter values, to classify the unmodified `CIFAR-10` training set and tested each model instance on the two *o.o.d.* generalisation test sets. We trained the networks with every combination of constraints — using either ReLU or Sigmoid unit activation, with or without a bottleneck, while varying the level of internal noise. The results were averaged over four random seeds for each network.

### 4.2. Results and Discussion

Figure 7 shows the change in generalisation accuracy for `small-inception` networks, trained with a particular value of internal noise, relative to a baseline network trained without activation noise.^10^ The inset shows the change in accuracy from the baseline model. Consistent with previous findings (Geirhos et al., 2018), the networks are not much affected by the very low levels of image degradation, but increasing it even slightly further rapidly lowers the accuracy of all models to chance performance (10%). In contrast, human performance decreases much more gradually with increased degradation (Geirhos et al., 2018).

**Figure 7:**
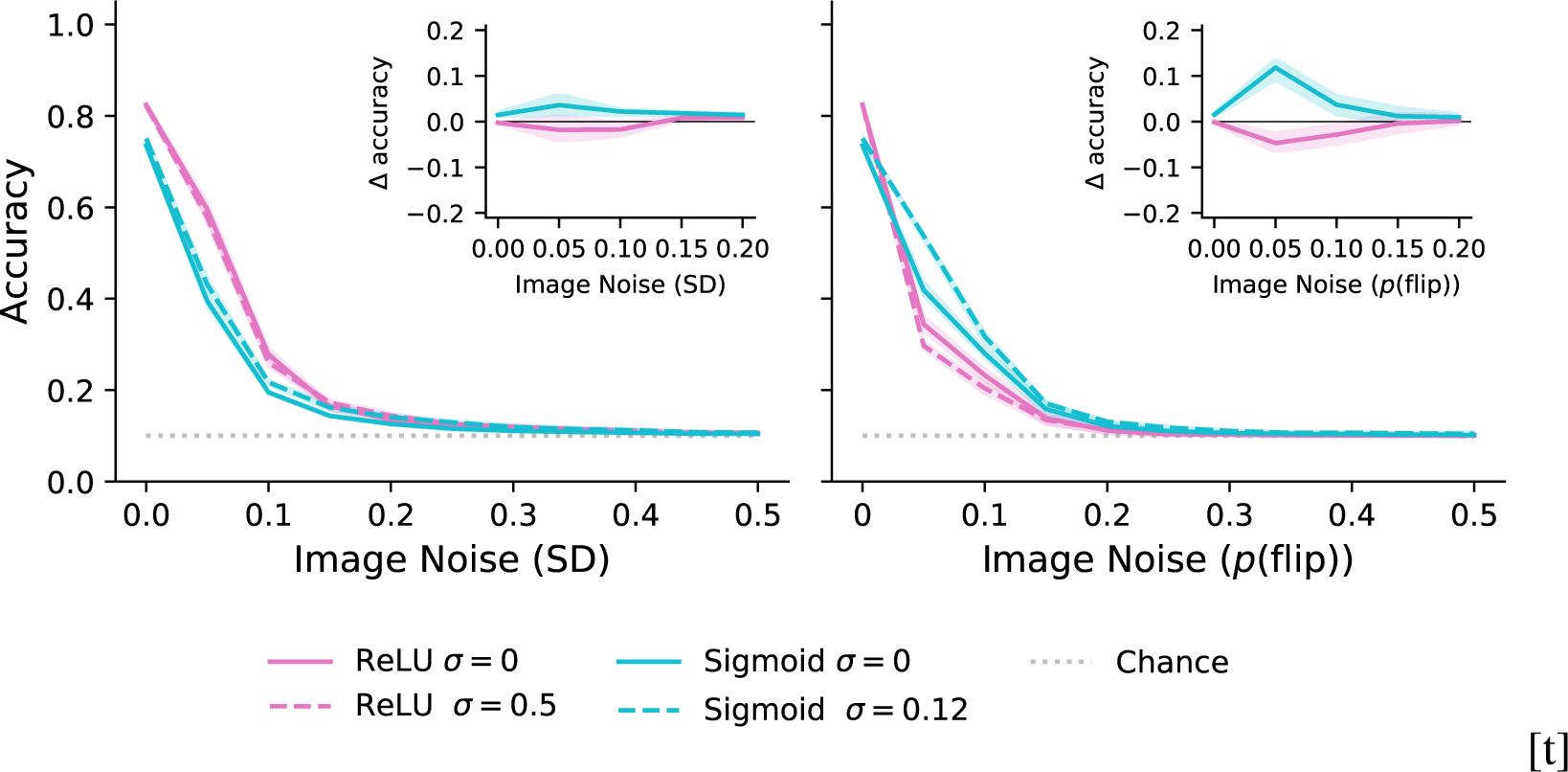
small`-inception` networks with internal noise have better generalisation for low levels of input degradation for Sigmoid (teal), but not ReLU (pink) activation function. **Left**: Uniform noise. **Right**: Salt-and-pepper noise. Solid lines represent baseline models trained without constraints. Dashed lines represent networks trained with one specific noise level (0.5 for ReLU and 0.12 for Sigmoid). Insets highlight deviation of models with internal noise from the baseline model (distance from the midline). So values above the *x*-axis show a better performance compared to baseline while values below show a worse performance.

For the networks with ReLU activation, the accuracy of the models with internal noise was 5 − 10% lower than the performance of the baseline models, for both additive uniform and salt-and-pepper degradation. That is, training with noise offered no advantage in generalisation to these images. On the other hand, the CNNs using sigmoid activations and internal noise did show improvement over the baseline model for both types of degradation (see Insets in Figure 7). The size of the effect was small for uniform noise (< 5%) and somewhat larger for salt-and-pepper noise (∼ 10%). The magnitude of the effect was also enhanced by the addition of a bottleneck (Appendix C, Figure C.2). However, this advantage was observed only for low levels of image degradation and quickly disappeared as the performance of all models deteriorated. Furthermore, the advantage was either not observed, or considerably smaller for `small-alexnet` networks (see Appendix A, Figure A.2).

The constraints appear to have slightly improved generalisation abilities for some combinations of conditions, but not for others, and only for some low levels of noise. As the advantage is further contingent on factors such as the model architecture, we conclude that the results do not support our prediction that training with constraints will overall improve robustness to image distortions. Clearly, additional constraints are needed to explain the human capacity to generalise to *o.o.d.* datasets. All models, both with and without constraints, quickly deteriorated as the amount of image degradation increased, at a similar pace. Only the networks with Sigmoid activation showed an improvement in accuracy, but only for low amounts of input noise. Furthermore, it is not clear why the effect is larger for salt-and-pepper noise compared with uniform noise. Finally, it should be taken into account that in absolute terms the accuracy of ReLU networks for uniform input noise was consistently higher than that of Sigmoid networks. For salt-and-pepper noise, the reverse was true — CNNs with sigmoid activations performed better than networks with ReLU units.

## 5. General Discussion

If CNNs are to be considered models of human vision they should not only succeed in classifying images under conditions in which humans succeed, but also fail under conditions in which human fail. Here we carried out a series of experiments that explore a striking example of CNNs outperforming humans, namely, in their ability to classify random data, in the form of images that look like TV static (random pixel images) or naturalistic images that are randomly assigned to categories (shuffled labels), as reported by Zhang et al. (2017). Our key contribution is to introduce biological constraints to CNNs that allow them perform well on unmodified naturalistic image data but fail on random data. These findings suggest that these constraints, or some version of them, should be implemented in CNNs when modelling human vision in order to provide better alignment between their performance characteristics.

To summarise, we report five main findings. First, in two behavioural experiments we confirm that humans perform far worse when learning random data compared to standard CNNs (Figure 2). In addition, we observed that humans are able to learn (a small set of) images with randomly assigned labels, but are mostly unable to learn images of random pixels. This behaviour contrasted with CNNs, which learn random pixels either more easily Zhang et al. (2017), or as in our simulations, learn the two types of random data at similar rates, with random pixels slightly easier than shuffled labels in some conditions and slightly harder in others.

Second, we found that exposing CNNs to structured data (pre-training on `ImageNet`) did not prevent them from learning random data. Surprisingly, pre-training actually made it easier for CNNs to learn to classify the random datasets (Figure 3), suggesting that this superhuman capacity reflects their architectures rather than their background training history.

Third, we introduced three biologically inspired modifications to CNNs designed to reduce their resource capacities. The addition of activation noise designed to model neural noise was the most important factor in selectively reducing performance on random data, but the introduction of a bottleneck (modelling the optic nerve) and a sigmoidal activation function (corresponding to the bounded firing rate of neurons) contributed to more human-like performance as well (Figure 5, **Top**).

Fourth, we show that these biological constraints worked by reducing network capacity rather than through some alternative mechanism. That is, we showed that CNNs with these constraints could still learn noise patterns, but far fewer of them (Figure 5, **Bottom**). This is the pattern of results to be expected if the manipulation interferes with capacity.

Finally, we did not find that the pre-training and the biological constraints helped much to improve better generalisation to *o.o.d.* data. Indeed, most of the networks did not improve on, or fared worse than baseline models with no constraints when tasked to classify test set images perturbed with different types of noise (Figure 7). Constraints did appear to improve performance slightly in CNNs with sigmoid activations, but benefits were mostly restricted to low levels of one particular type of input noise.

What do our results say about the correspondence between CNNs and human vision? The first point to note is that our findings highlight the importance of adding biological constraints that reduce CNN resource capacity when attempting to model human vision. Although the extraordinary success of state-of-the-art CNNs in image classification is often taken as evidence that CNNs are promising models of human vision, these models are clearly exploiting resource capacities that far exceed those available to networks in the human visual system, allowing models to learn random data almost as well as structured data. Biologically plausible CNNs need to succeed in classifying images without the ability to learn random data and our own findings highlight three constraints that appear to be relevant to this.

However, clearly the pre-training and the biological constraints are not enough to reconcile the CNN and human results. First, the networks do not reproduce the pattern of relative difficulty of the random datasets that humans exhibit. Learning to classify random pixel images is much more difficult for humans than classifying structured objects in arbitrary categories. This pattern of results was not consistently observed with CNNs. And more importantly, the networks with capacity constraints, that were selectively good at classifying structured data, were not much better at *o.o.d.* generalisation. Clearly a range of additional constraints and perhaps new models are required to capture this key feature of human object recognition.

Arpit et al. (2017) have also explored how to reduce the memorisation of random data in CNNs, although they did not consider the relevance of their findings to human vision. Specifically, they assessed the ability of several commonly used regularisation techniques to reduce memorisation of random data. Interestingly, some of the findings of Arpit et al. (2017) differ from our own results. Most relevant to our work, they concluded that the introduction of internal Gaussian noise was ineffective as it reduced performance on both structured and unstructured data. By contrast, our results show that it is possible to find levels of noise that selective impair performance on unstructured data. It is not clear why we reached different conclusions, but it may be explained by differences in the implementation of models. Firstly, we applied internal noise both during training and testing of the networks. This is not conventional practice when using regularisers, but bears closer correspondence to the biological brain. Secondly, Arpit et al. (2017) do not describe the methodological details of their procedure, such as at which points within a network the noise is added. This lack of procedural details prevents further comparisons between the results of the two studies.

It is interesting to note that a similar pattern of learning unstructured data is observed in the domain of natural language processing as well. That is, while deep networks show impressive successes in learning natural languages, unlike humans, they also find it easy to learn impossible languages (Mitchell & Bowers, 2020). Both findings highlight missing inductive biases in standard CNNs that result in networks with superhuman learning capacities in some domains. A key challenge in creating neural network models with greater correspondence to human performance is to introduce the relevant constraint and biases so that models learn what they should learn (under realistic training conditions), and at the same time do not learn what humans cannot.

In summary, we have found that adding three basic biological constraints to CNNs leads to them exhibiting more human-like performance through impeding their ability to classify random data, while preserving their ability to classify structured data. However, models trained with the examined constraints are not better at *o.o.d.* generalisation. Developing models that succeed where humans succeed, but also fail where humans fail, is an important challenge for future work in developing better models of human vision.

## Acknowledgements

This project has received funding from the European Research Council (ERC) under the European Union’s Horizon 2020 research and innovation programme (Grant Agreement No. 741134).

## Appendix A.

This section contains figures displaying results for several simulations described in the Section 3 for networks with the `small-alexnet` model architecture. Results from Figure A.1 correspond to those for `small-inception` from Figure 5. Figure A.2 corresponds to Figure 7.

**Figure A.1:**
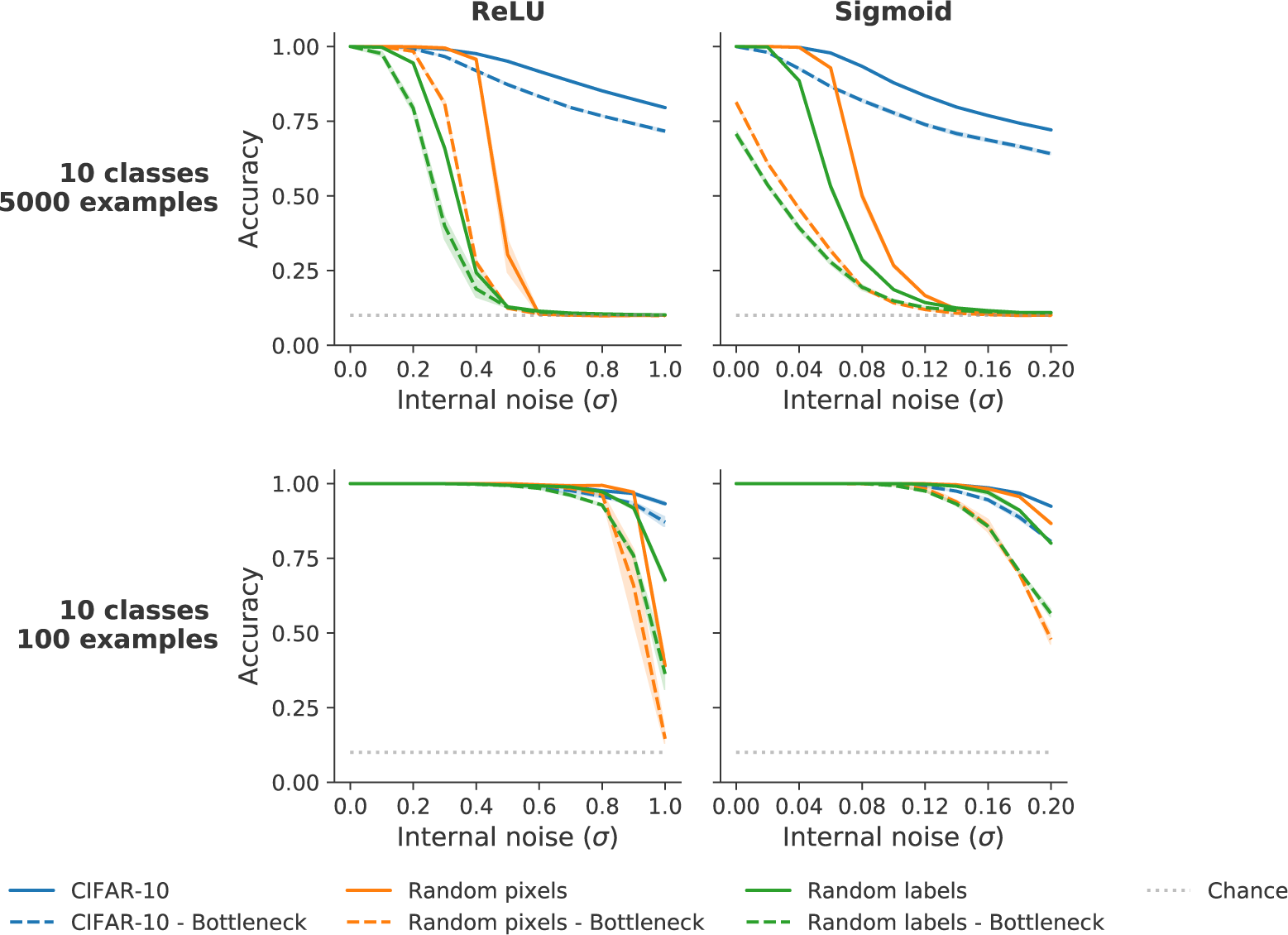
**Top**: Accuracy for `small-alexnet` models with biological constraints. **Left**: ReLU activation function. **Right**: Sigmoid activation function. The graphs show the impact of the capacity manipulations on training accuracy for the three types of datasets (blue: unmodified `CIFAR-10`; green: modified `CIFAR-10` with shuffled labels; orange: dataset of random pixels images). The internal noise levels used are plotted on the *x*-axis. Solid lines indicate model accuracy for models without a bottleneck; dashed lines indicate models with a bottleneck. **Bottom**: Training set accuracy for models trained with biologically inspired constraints. Networks trained on `CIFAR-10` images, with either unmodified (**blue**) or shuffled (**green**) labels, and random pixel images (**orange**). The CNNs architecture either included a bottleneck (**dashed lines**), or did not (**solid lines**). Hidden units used either ReLU (**left**) or Sigmoid (**right**) activation functions. Networks trained with a limited number of examples per class (10 classes with 100 examples).

**Figure A.2:**
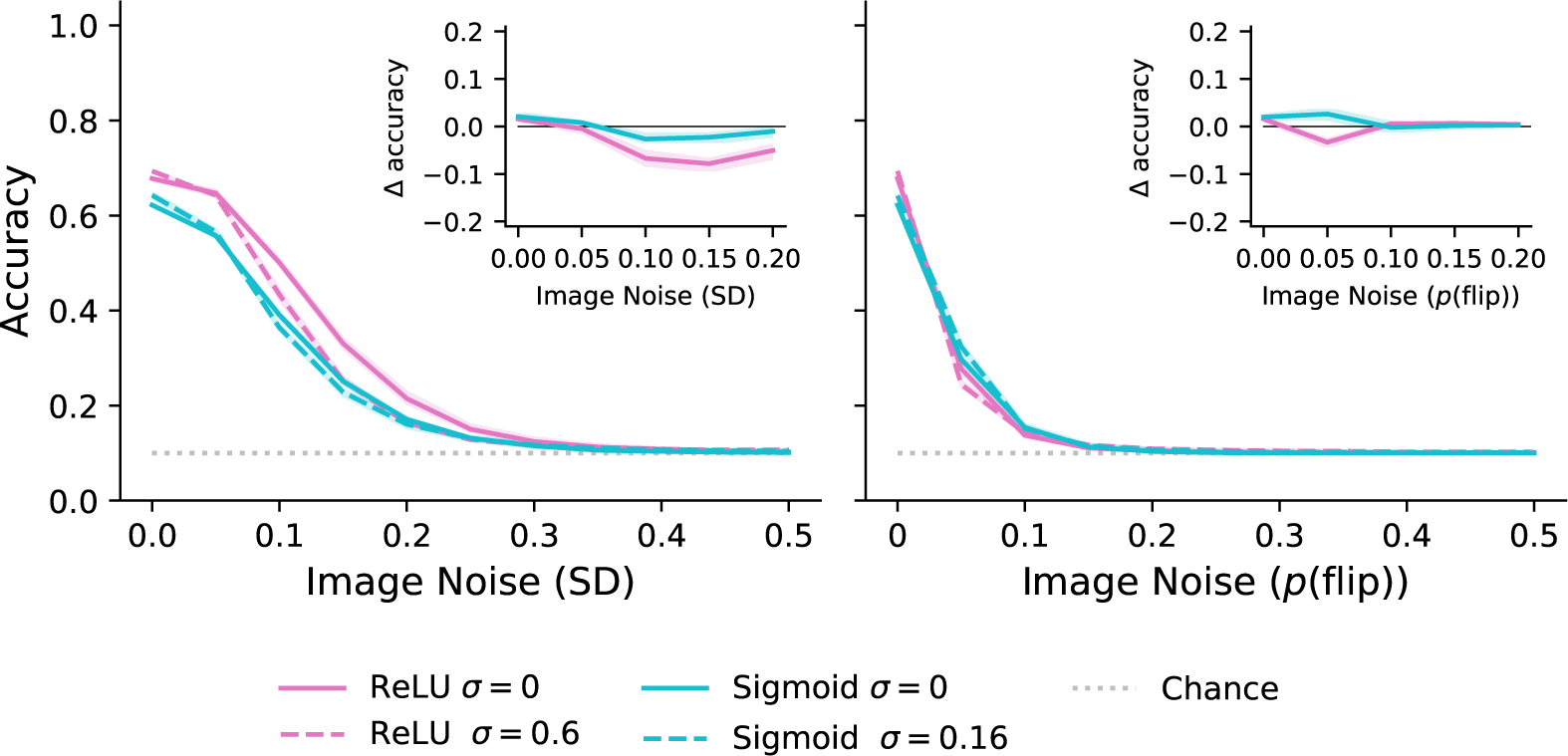
`small-alexnet` models trained with internal noise (orange lines) do not generalise to novel distorted images better than a baseline model (blue lines). **Left**: Uniform noise. **Right**: Salt-and-pepper noise. Solid lines represent baseline models trained without constraints. Dashed lines represent networks trained with one specific noise level (0.6 for ReLU and 0.16 for Sigmoid). Insets highlight deviation of models with internal noise from the baseline model (distance from the midline). Values above the *x*-axis show a better performance compared to baseline while values below show a worse performance.

## Appendix B.

Neural networks that classify random data are often described as performing ‘memorisation’ (Zhang et al., 2017; Arpit et al., 2017). The basic idea is that networks are representing each specific input pattern rather than extracting common features amongst members of a given class. Indeed, that must be the case given that there are no common shared features of members of a given class, as with random pixel images. One plausible prediction is that CNNs that classify images on the basis of memorising all the training patterns will not only fail to generalise well to test stimuli, but will also generalise less effectively if the training data were to be perturbed in a way not experienced during training. Indeed, learning invariant patterns is often thought to support generalisation in CNNs.

To explore generalisation in CNNs trained to memorise random data we trained `small-alexnet` networks to classify random pixel images. Afterwards, taking a smaller sample of the training dataset, we changed the values of a randomly selected proportion of the pixels in each image. The new values were sampled from a uniform distribution *a ∼ U* (0., 1.). We then assess the accuracy of the networks on this modified training data. We find that, while network performance decreases as a larger proportion of the original pixel values are changed, their accuracy remains relatively high (Figure B.1). This effect is present in both ReLU (pink) and Sigmoid (teal) networks.This highlights that CNNs trained on random data can generalise in the sense that they are robust to perturbations in the data they were trained on.

**Figure B.1.**
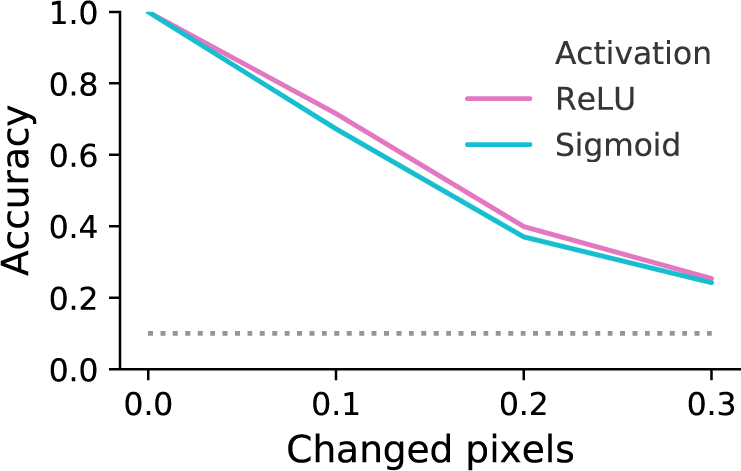
: Prediction accuracy of `small-alexnet` models with ReLU (pink) or Sigmoid (teal) activation on modified random pixel training data. The CNNs performance degrades gradually as a greater proportion of pixels is changed, not all at once.

## Appendix C.

The figures below show the generalisation performance of CNNs trained with internal noise plus a bottleneck, for ReLU and Sigmoid units. They correspond to the results depicted in Figures 7 and A.2.

**Figure C.1:**
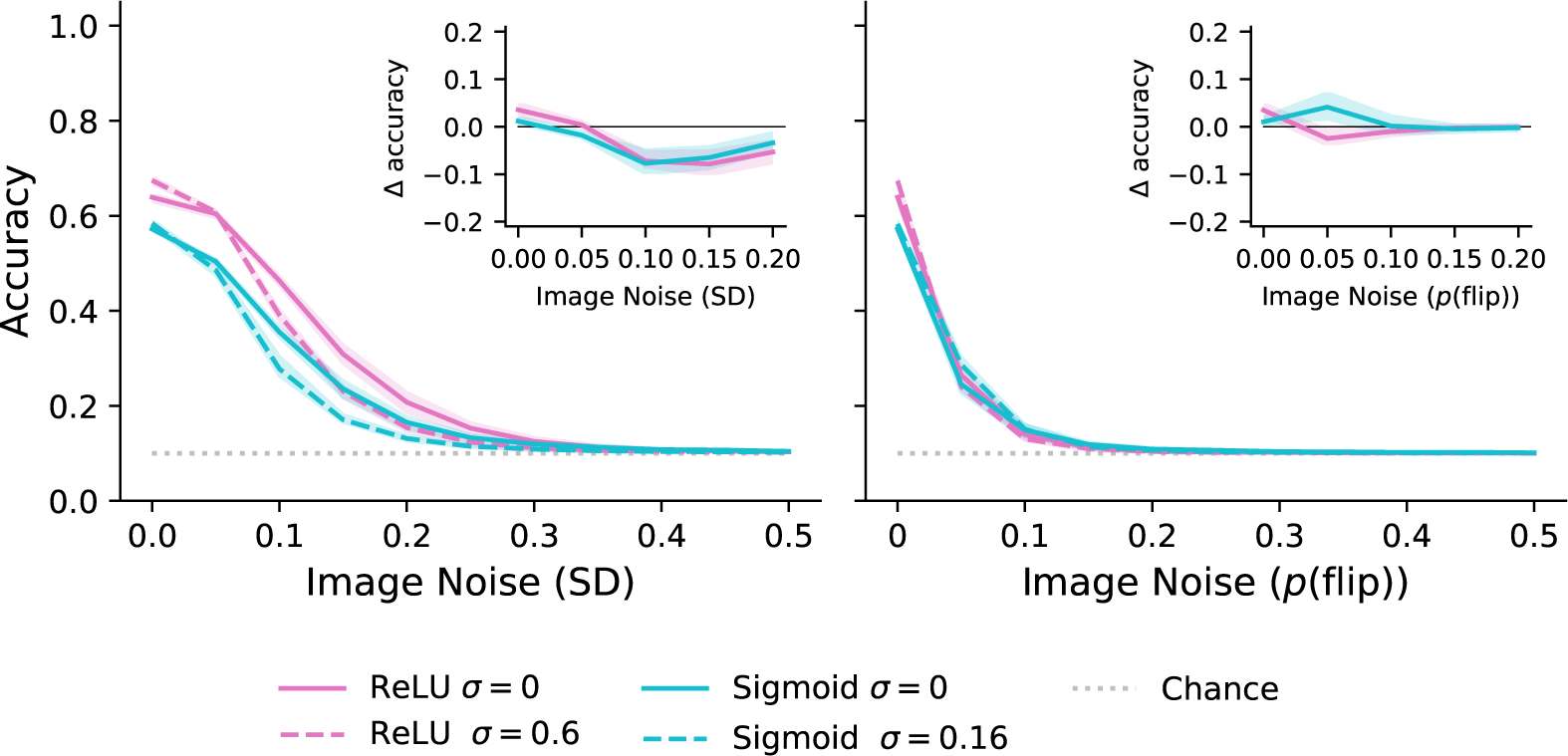
`small-alexnet` models trained with internal noise and a bottleneck (dashed lines) do not generalise to novel distorted images better than a baseline model (solid lines). **Left**: both networks with ReLU (pink) and Sigmoid (teal) activations perform worse than the baseline on uniform input noise. **Right**: There is a slight improvement for Sigmoid networks with constraints (teal), but not ReLU (pink) over the baseline performance for salt-and-pepper input noise.

**Figure C.2:**
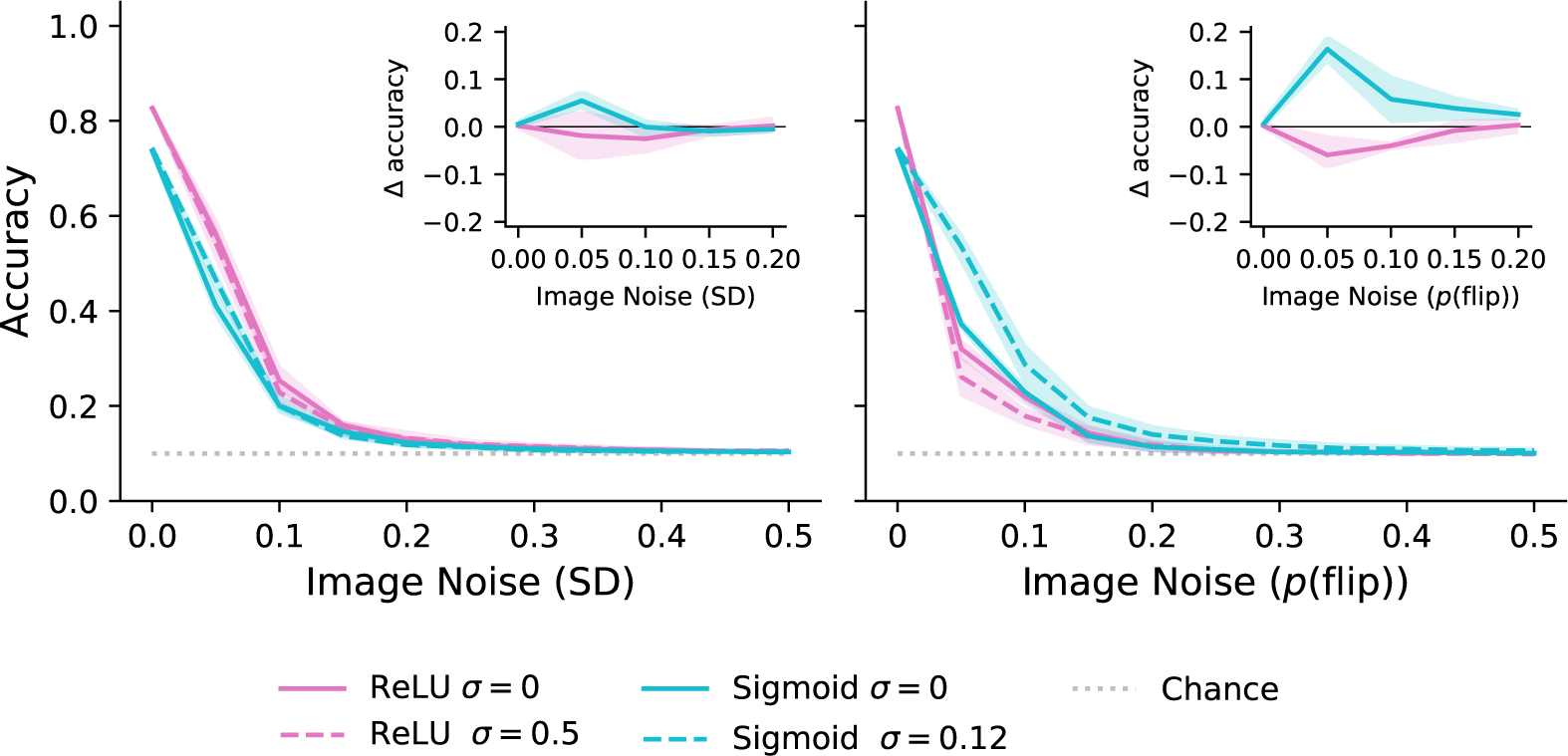
`small-inception` models trained with internal noise and a bottleneck (dashed lines) showed improved generalisation to novel distorted images over the baseline models (solid lines) for networks with Sigmoid units (teal), but not ReLU (pink). The effect was larger for images with salt and pepper noise (**right**) than for images with uniform noise (**left**). The average deviation in accuracy from the baseline model is highlighted in the inset, with values above the mid-line indicating an improvement in performance.

## Appendix D.

In this section we address the concern about whether insufficient training can explain our results from Section 3.2, as training with high levels of internal noise slows down learning. We replicated the simulations, doubling the number of training epochs to 200. We show that the additional training does not change the patter of results observed previously. Note that models with no internal noise were omitted as they do not pertain to the replication problem, and fewer values of internal noise were sampled.

**Figure D.1:**
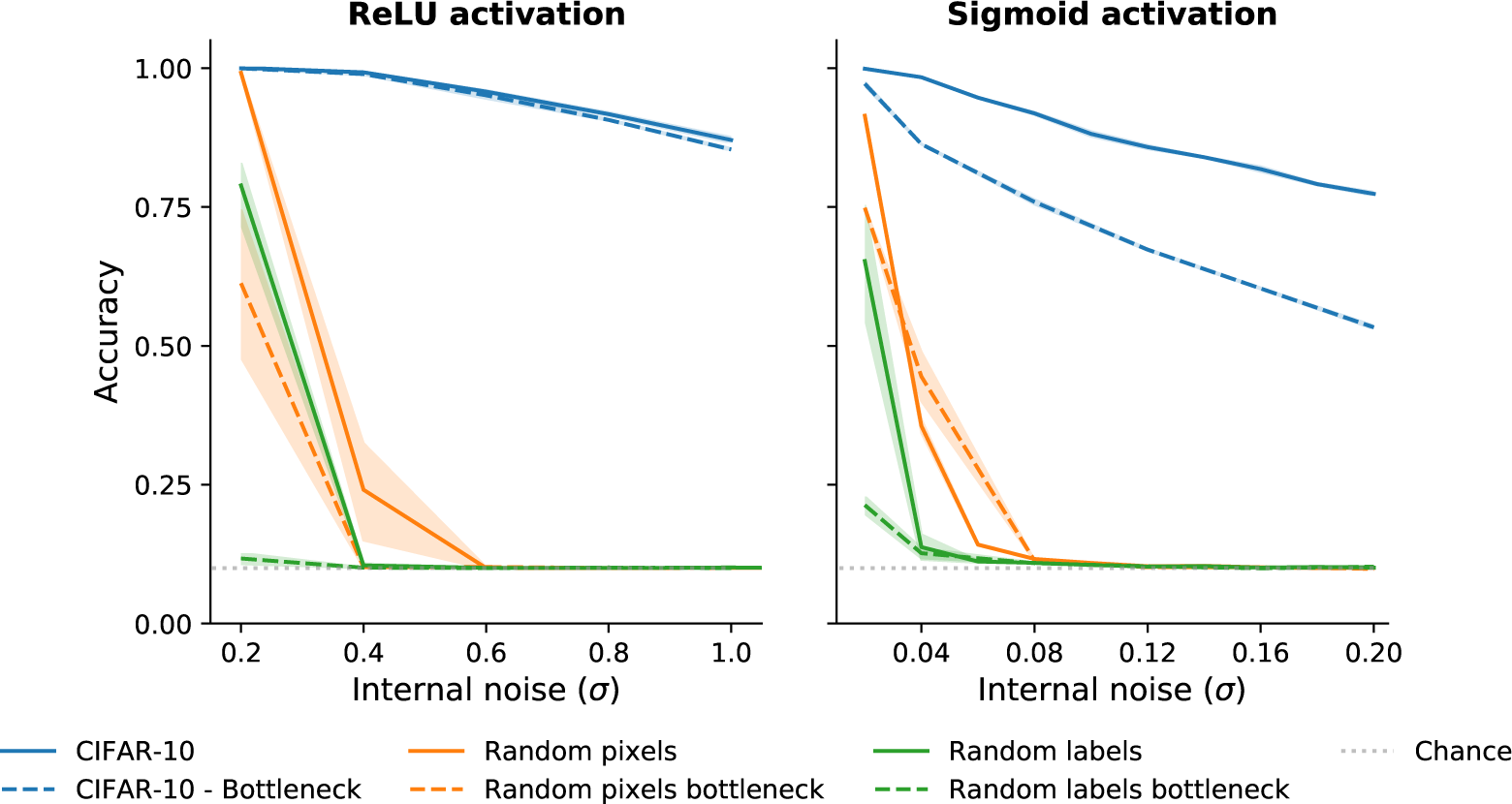
Accuracy for `small-inception` models with biological constraint trained. Replication of results from Figure 5-Top with 100 more epochs of training (total 200). The results are consistent with previous findings, showing the random pixels and shuffled labels datasets are not learned with after the extra training.

## Appendix E.

In this section we include results from two more architectures, `DenseNet` (Huang et al., 2016) and `MobileNet` (Howard et al., 2017), replicating the simulations in Section 3.2. Two different random seeds were used for each model instance for both networks. Results are analogous to those from Figures 5 and A.1 and replicate the qualitative relations between the different data types.

**Figure E.1:**
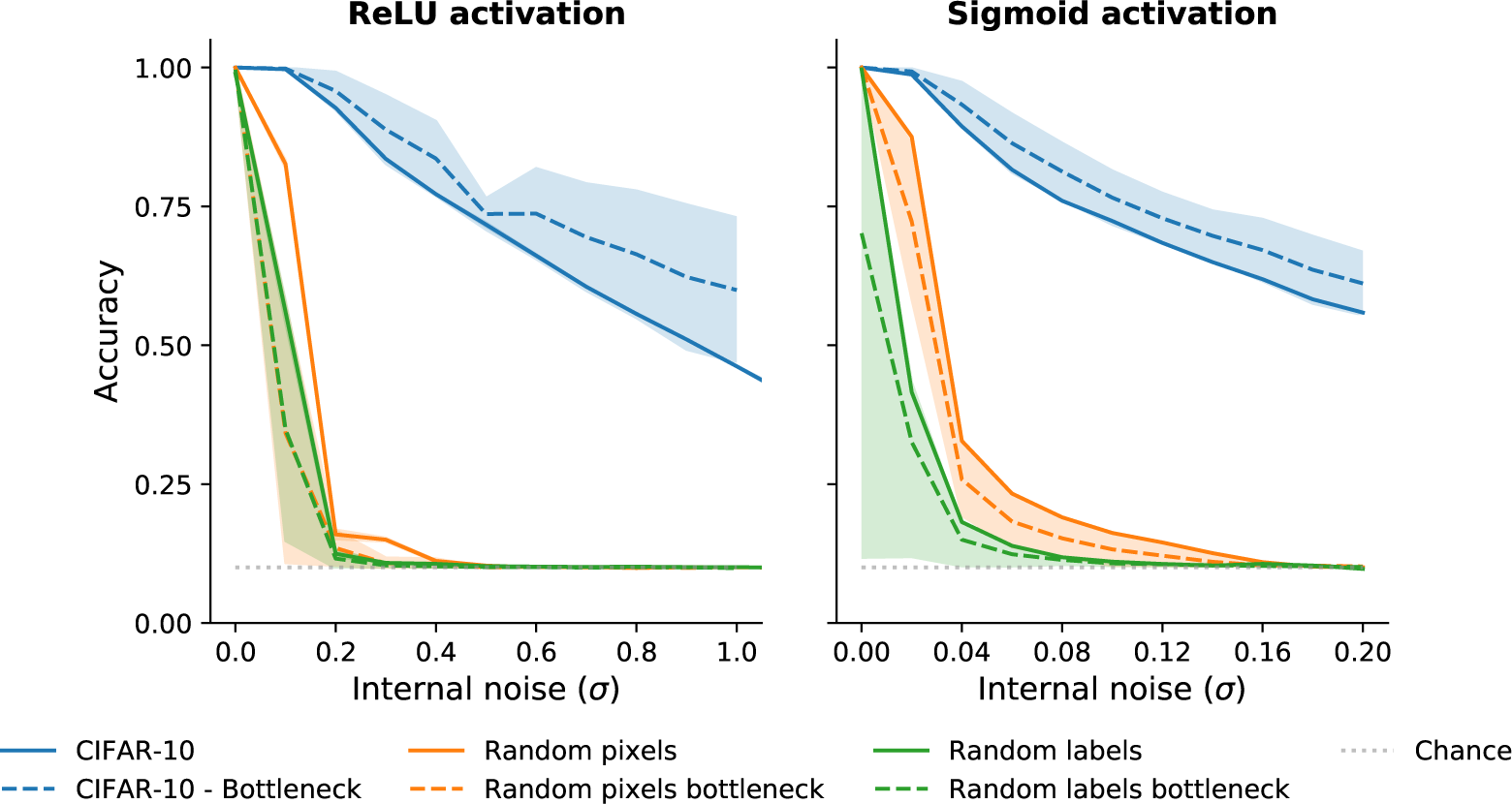
Accuracy for `DenseNet` models with biological constraints. **Left**: ReLU activation function. **Right**: Sigmoid activation function. Results are consistent with the pattern seen in Figures 5 (Top) and A.1. Interesting to note is the effect of the bottleneck appears to be more substantial for this architecture.

**Figure E.2:**
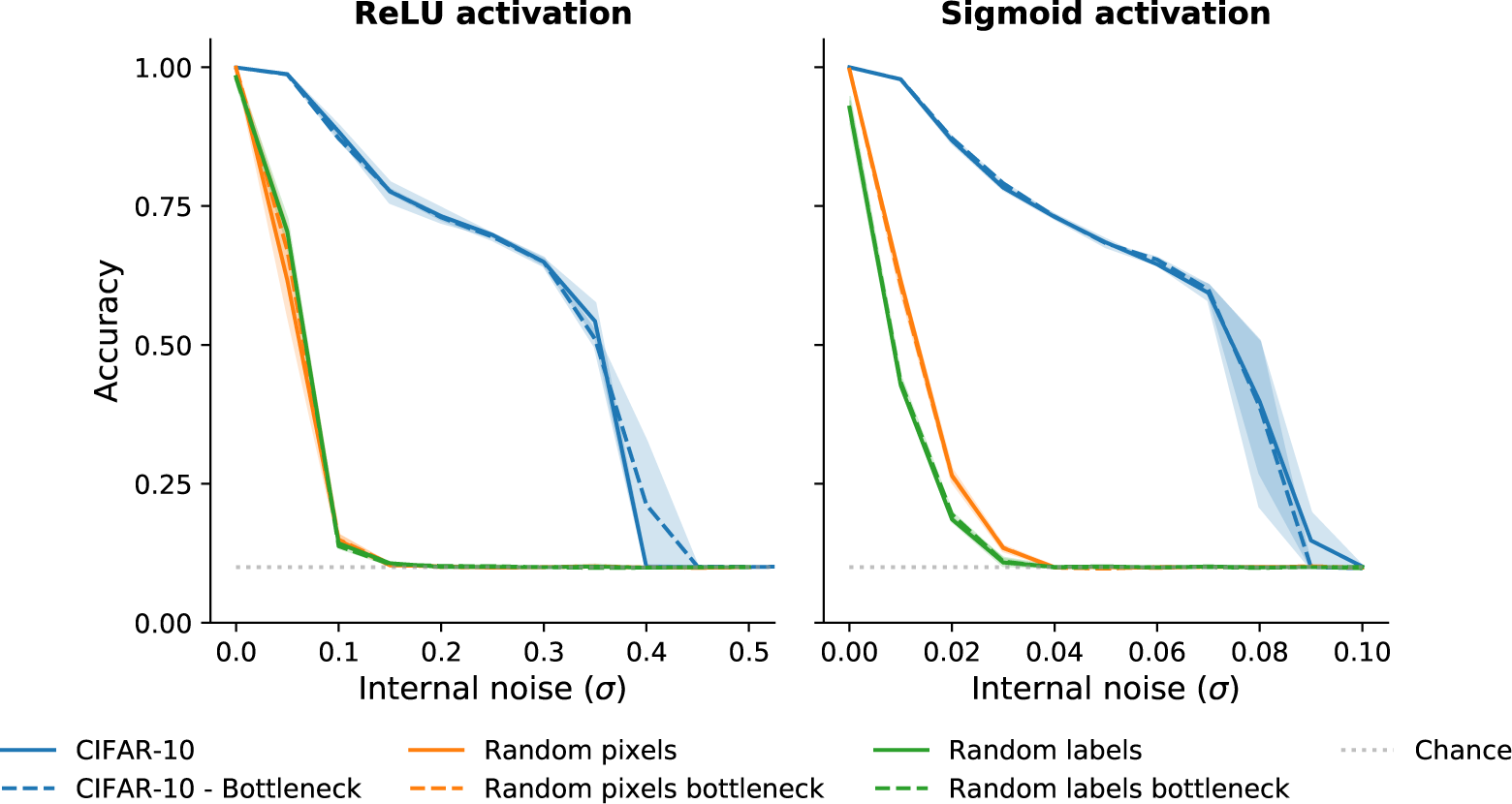
Accuracy for `MobileNet` models with biological constraints. **Left**: ReLU activation function. **Right**: Sigmoid activation function. Results are consistent with other simulations with capacity constraints.

## Appendix F.

In this section we provide a summary of the hyperparameters involved in training every model for all combinations of constraints. All models are trained using a standard Stochastic Gradient Descent (SGD) optimiser. The hyperparameter values declared for simulations in Section 3.2 were also used for all models and simulations in Section 4.

**Table F.1:** Hyperparameters for all models trained in the simulations.

| Model | Section | Batch | Epochs | LR | Decay | Momentum |
| --- | --- | --- | --- | --- | --- | --- |
| small-inception | 3.2 | 128 | 100 | $1 \times 10^{-1}$ | 0.05 | 0.9 |
| | Appendix D | 128 | 200 | $1 \times 10^{-1}$ | 0.05 | 0.9 |
| | 3.3 | 32 | 503 | $1 \times 10^{-1}$ | 0.05 | 0.9 |
| small-alexnet | Appendix A | 128 | 100 | $1 \times 10^{-2}$ | 0.05 | 0.9 |
| | Appendix A | 32 | 503 | $1 \times 10^{-2}$ | 0.05 | 0.9 |
| ResNet50 | 3.1 | 25 | 70 | $5 \times 10^{-4}$ | $1 \times 10^{-6}$ | 0.0 |
| DenseNet192 | Appendix E | 128 | 100 | $1 \times 10^{-2}$ | 0.05 | 0.9 |
| MobileNet | Appendix E | 128 | 100 | $1 \times 10^{-2}$ | 0.05 | 0.9 |

## Appendix G.

Here we present results from the experiments used to determine the bottleneck size suitable for each model presented in Section 3.2. The models were trained with bottlenecks of various sizes, defined as the number of feature maps in the first convolutional layer of the model. Results from these networks were compared with those from the baseline models without a bottleneck. The selection criterion was to choose the lowest number of feature maps that did not have a dramatically detrimental effect on the networks’ accuracy on the validation set compared to the baseline. All networks were trained to convergence, using three random seeds for each bottleneck size and the baselines. Since it was our goal to keep as many parameters constant as possible when comparing across conditions, it was desirable to use the same bottleneck size for networks with both ReLU and Sigmoid activation functions. We used the same hyperparameters as in Section 3.2, outlined in Appendix F.

For the `small-inception` networks using ReLU activations (Figure G.1, Left) lowering the number of feature maps down to 2 has a very small effect on the networks’ validation accuracy compared to the baseline model. However, a larger decline can be seen in the networks with 1 feature map. An overall sharper decrease can be observed for the networks with Sigmoid activations (Figure G.1, Right), where there is also a more pronounced drop-off from 2 to 1 feature map. We therefore decided to use a bottleneck of 2 feature maps for all further `small-inception` simulations.

We used a similar method to establish the size of bottleneck for the `small-alexnet` network. Here we noticed that validation accuracy dropped slowly yet steadily when we decreased the size of bottleneck (Figure G.2). The decline in accuracy was steeper for bottlenecks with fewer than 16 feature maps for both the networks using ReLU (left) as well as those using Sigmoid units (right). At this point there was a trade-off between the validation accuracy, which we wanted to remain high, and bottleneck size, which we wanted to be low to constrain the capacity of the network. Therefore, we selected a bottleneck size of 8 feature maps for simulations involving `small-alexnet` networks as the best middle-ground solution.

**Figure G.1:**
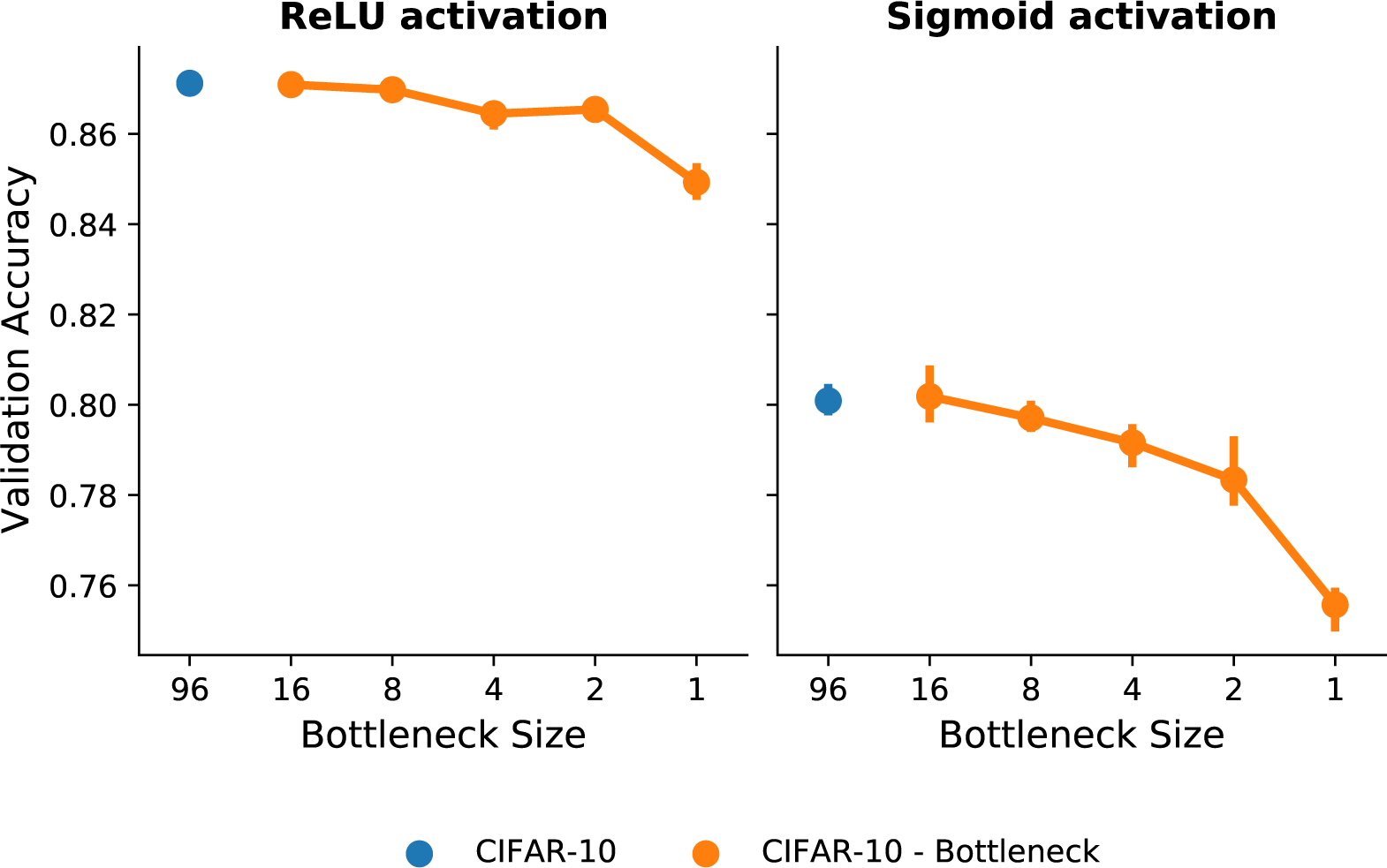
Validation accuracy for different bottleneck sizes for the `small-inception` architecture. Note the values on the *x*-axis represent levels of a categorical variable. Models with bottlenecks (orange) were compared with a baseline model with no bottleneck (blue). The networks used either ReLU (**Left**) or Sigmoid (**Right**) activation functions. Models with ReLU activations perform well on the validation set compared to the baseline model for bottleneck sizes as low as 2 feature maps. There is a steeper decline in accuracy between bottlenecks of 2 and 1 feature maps. This effect is even larger for networks using Sigmoid activations. Therefore 2 feature maps was selected as the optimal bottleneck size.

**Figure G.2:**
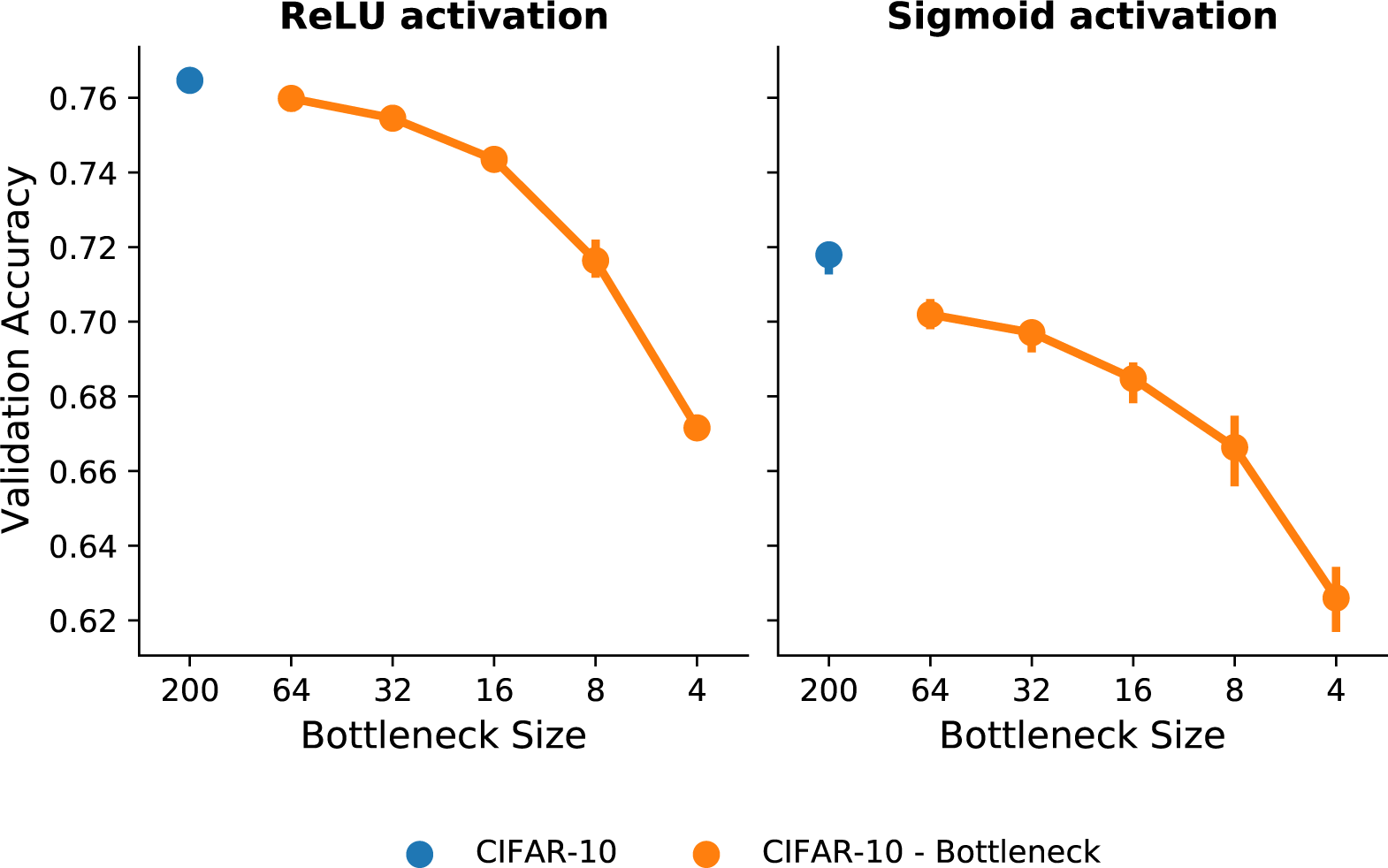
Validation accuracy for different bottleneck sizes for the `small-alexnet` architecture, for networks using either ReLU (**Left**) or Sigmoid (**Right**) activation functions. Note the values on the *x*-axis represent levels of a categorical variable. Models with bottlenecks (orange) displayed gradually lower validation accuracy compared to the baseline model (blue). The sharpest drop in validation accuracy was between 8 and 4 feature maps. Therefore, 8 feature maps was chosen as the preferred bottleneck size.

## Footnotes

1 Learning random datasets was characterised as memorisation (Zhang, et al., 2017; Arpit et al., 2017), but we note that, while CNNs trained on these datasets cannot generalise to entirely new examples, they do generalise to perturbed exemplars of studied items. See Appendix B.

2 Research code for all simulations and analyses is available at https://github.com/chris7sv/capacity_constraints

3 Note that, unlike the findings of Zhang, et al. (2017), the networks trained on the shuffled labels dataset converged faster than those trained on random pixel images. We found that this relationship was not a robust one, and varied based on e.g., the choice of architecture, or even the learning rate.

4 We have successfully replicated the results of these simulations with DenseNet (Huang, et al., 2016) and MobileNet (Howard et al., 2017), see Appendix E.

5 Implemented as a regularisation layer from tensorflow.keras module.

6 See Appendix G for results from these experiments.

7 To confirm our results were not due to insufficient training, we repeated all simulations for 200 epochs. Results are identical with those described in this section (see Appendix D).

8 Results for small-alexnet are qualitatively similar — see Appendix A.

9 See Appendix A, Figure A.1 for comparable results for small-alexnet models.

10 See Appendix C for results for models with bottleneck.

